# Robust spatial ventriloquism effect and aftereffect under memory interference

**DOI:** 10.1101/2020.09.02.279323

**Authors:** Hame Park, Christoph Kayser

**Affiliations:** Department for Cognitive Neuroscience, Faculty of Biology, Bielefeld University, Universitätsstr. 25, 33615, Bielefeld, Germany; Center for Cognitive Interaction Technology (CITEC), Bielefeld University, Inspiration 1, 33615, Bielefeld, Germany

**Keywords:** multisensory integration, multisensory recalibration, memory, audio-visual, spatial perception

## Abstract

Our brain adapts to discrepancies in the sensory inputs. One example is provided by the ventriloquism effect, experienced when the sight and sound of an object are displaced. Here the discrepant multisensory stimuli not only result in a biased localization of the sound, but also recalibrate the perception of subsequent unisensory acoustic information in the so-called ventriloquism aftereffect. This aftereffect has been linked to memory-related processes based on its parallels to general sequential effects in perceptual decision making experiments and insights obtained in neuroimaging studies. For example, we have recently implied memory-related medial parietal regions in the trial-by-trial ventriloquism aftereffect. Here, we tested the hypothesis that the ventriloquism aftereffect is indeed susceptible to manipulations interfering with working memory. Across three experiments we systematically manipulated the temporal delays between stimuli and response for either the ventriloquism or the aftereffect trials, or added a sensory-motor masking trial in between. Our data reveal no significant impact of either of these manipulations on the aftereffect, suggesting that the recalibration reflected by the ventriloquism aftereffect is surprisingly resilient to manipulations interfering with memory-related processes.

## INTRODUCTION

Sensory recalibration is a mechanism by which the brain continuously adapts to apparent discrepancies in our sensory environment, such as the displaced figure and voice of an actor in a movie watched over headphones^1,2^. One example of adaptive recalibration is the ventriloquism aftereffect, a frequently studied paradigm for multisensory perception in the laboratory. To reveal this aftereffect, participants are first (in an audio-visual trial) presented with spatially discrepant audio-visual stimuli, which give rise to a biased localization of the sound. This bias reflects the partial fusion of the discrepant audio-visual information – the so called ventriloquism effect^3^. In a subsequent trial, participants are then asked to localize a unisensory sound, which they often misjudge in the direction established by the previous multisensory discrepancy^4–7^. For example, when the light is to the left of the sound in the audio-visual trial, the subsequent sound is misjudged to the left. This aftereffect, or recalibration bias, is observed following prolonged exposure to consistent multisensory discrepancies^8,9^, but also following single trial exposure, the so called immediate or trial-by-trial recalibration effect^4,7^. In either case, the resulting aftereffect bias reflects the persistent influence of previously received multisensory evidence on subsequent behavior.

Using neuroimaging we have recently investigated the neurophysiological mechanisms underlying the trial-wise ventriloquism aftereffect^7^. We found that medial parietal cortices reflect the persistent encoding of previous multisensory stimuli and are predictive of the trial-wise aftereffect^7^. This led us to speculate that brain regions traditionally implied in spatial and working memory^10–13^ contribute to the ventriloquism aftereffect, for example by maintaining a representation of previous sensory evidence between trials and mediating its influence on the perception of subsequent stimuli. Such a role of parietal regions in the ventriloquism aftereffect has also been suggested by other studies, and possibly the same parietal processes contribute to both the immediate and long term effects^14,15^.

That brain regions involved in short-term memory may contribute to the ventriloquism aftereffect is similarly predicted by studies on other types of serial dependencies in perceptual decision making. Statistical dependencies between judgements made in consecutive trials are seen ubiquitously in sensory and cognitive tasks^16–18^. While in many paradigms such dependencies could in principle arise from changes in early sensory representations, the emerging consensus seems to be that these arise from memory-related processes^17,19^. In support of this, recent studies showed that experimental manipulations known to affect memory processes^20,21^, such as changing the delay between stimulus and response, alter serial dependencies during judgements of visual orientations^22^, the accumulation of the ventriloquism aftereffect^9^, and longer reaction times reduce perceptual bias in visual discrimination^23^. Collectively, the functional analogy of the trial-wise ventriloquism aftereffect with serial dependencies in perceptual decision making and the neuroimaging studies implying medial parietal regions in the aftereffect, make a strong case for a memory-related component in the ventriloquism aftereffect.

We set out to test the hypothesis that the trial-wise ventriloquism aftereffect is related to memory processes, and hence susceptible to manipulations known to interfere with working memory. In three experiments we i) manipulated the delay between the inducing audio-visual (ventriloquist) stimulus and the associated response, ii) manipulated the delay between stimulus and response in the auditory trial, or iii) used a masker trial in between the audio-visual and the auditory trial to interfere with mnemonic processes. We found that none of the manipulations led to a consistent and robust change in the aftereffect bias, suggesting that the ventriloquism aftereffect is more robust to memory-manipulations as expected from similar studies on serial dependencies in serial perception.

## RESULTS

### Multisensory response biases

In three experiments we probed participants’ judgments of sound location in audio-visual (AV) trials and subsequent auditory (A) trials (**Figure 1**). In the AV trials, spatially localized sounds were accompanied with spatially localized random-dot patterns presented at either the same location or a range of spatial discrepancies (ΔVA). This allowed us to quantify the ventriloquism effect, reflecting the bias induced by the visual stimulus on the perceived location of the simultaneous sound. The responses in the subsequent A trials allowed us to probe the trial-wise ventriloquism aftereffect, reflecting the persistent influence of the multisensory discrepancy experienced in the AV trial on the judgement of a subsequent unisensory sound^4,7^. Each experiment manipulated the sequence of AV-A trials in a different manner: experiment 1 induced a variable delay before the response in the AV trial, experiment 2 induced a variable delay before the response in the A trial, and experiment 3 introduced a sensory-motor masker stimulus in between AV and A trials (**Figure 1**).

**Figure 1.**
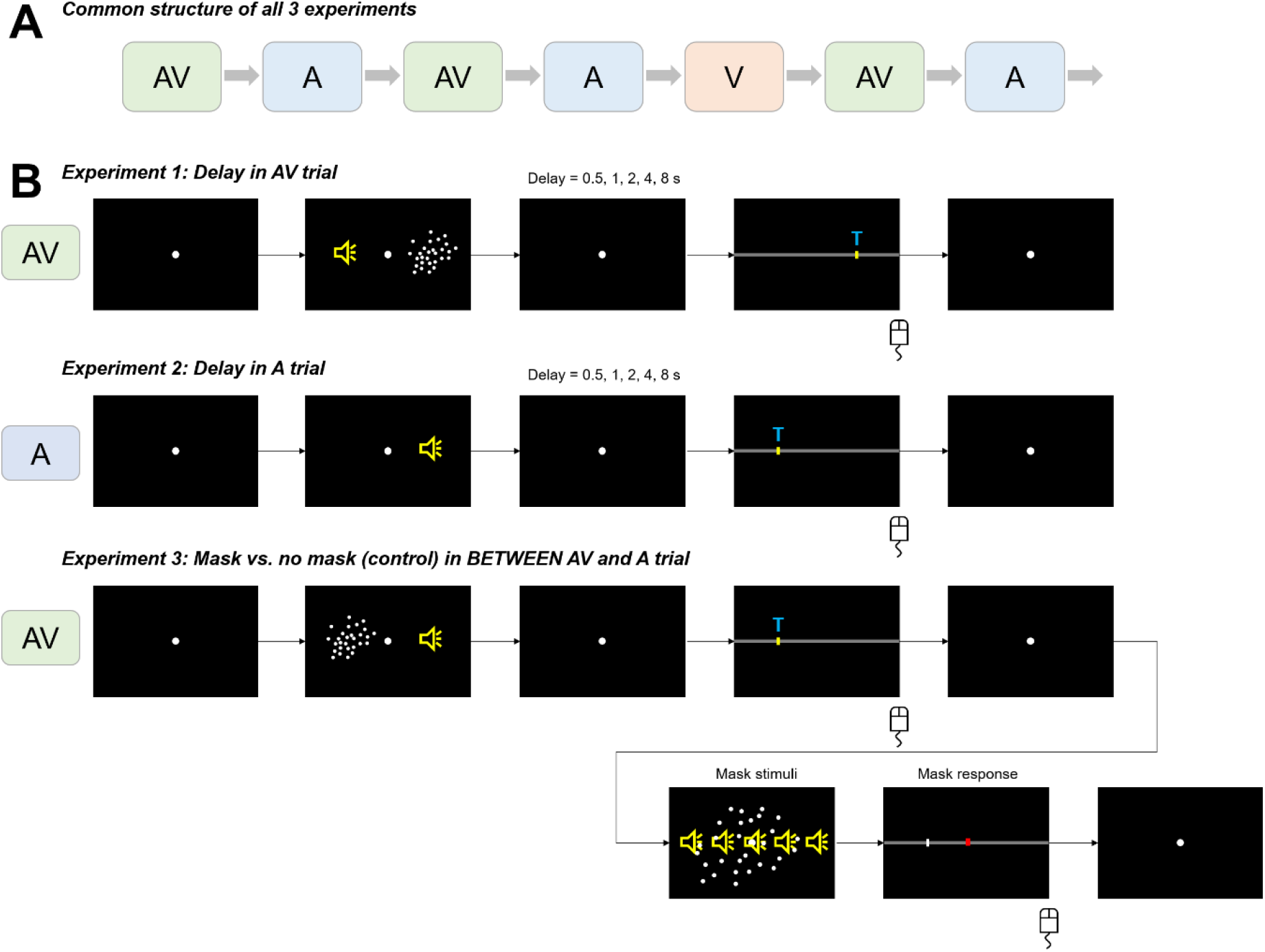
Experiment design and trials. **(A)** All 3 experiments consist of the same basic structure of interleaved AV-A trials and occasional V trials not interrupting the AV-A sequence. **(B)** The experimental manipulation of each experiment. The top sequence shows the AV trial with a varying delay between stimulus and response (Experiment 1). The middle sequence shows Experiment 2, in which the delay between stimulus and response in the A trial was manipulated. In both Experiments 1& 2 the inter-trial intervals had a default delay of 800 ms – 1200 ms (uniform). The bottom sequence shows the masking trials inserted in between the AV and A trials in Experiment 3. The masking trial was present in half the AV-A sequences, in the other half there was no masking trial (control) but the inter-trial interval between AV-A trials was extended (1800 ms – 2000 ms) to obtain an overall similar delay between AV and A trials in the masking and control conditions.

In a first step we determined the dependency of each bias on the multisensory discrepancy, ΔVA. For this we combined the data across all three experiments and, following previous studies^14,24,25^, compared linear and non-linear models in their predictive power for each bias (eq. 1-3). This revealed ‘very strong’ evidence that the ventriloquism bias was best explained by a combined linear and non-linear dependency on ΔVA: relative group-level BIC values = 87, 74, 0, for models 1-3 (c.f. Materials and Methods). In contrast, the aftereffect was best described as a linear dependency on ΔVA: rel-BIC = 0, 11, 11 (models 1-3). In the following we hence focused on the combined linear and nonlinear model (eq. 3) for the ventriloquism bias and a linear model (eq. 1) for the aftereffect to probe whether these are affected by the experimental manipulations.

### Manipulating the delay within audio-visual trials

In the first experiment (n=20) we manipulated the temporal delay between the audio-visual stimulus and participant’s response in the AV trial, which could take one of the five average values (0.5s, 1s, 2s, 4s, and 8s; ± 200 ms uniform random jitter in each trial). This manipulation could in principle affect both the ventriloquism bias and the aftereffect bias. **Figure 2A** shows the resulting biases (as participant-averaged data) for the two extreme values of the delay (0.5 and 8s).

**Figure 2.**
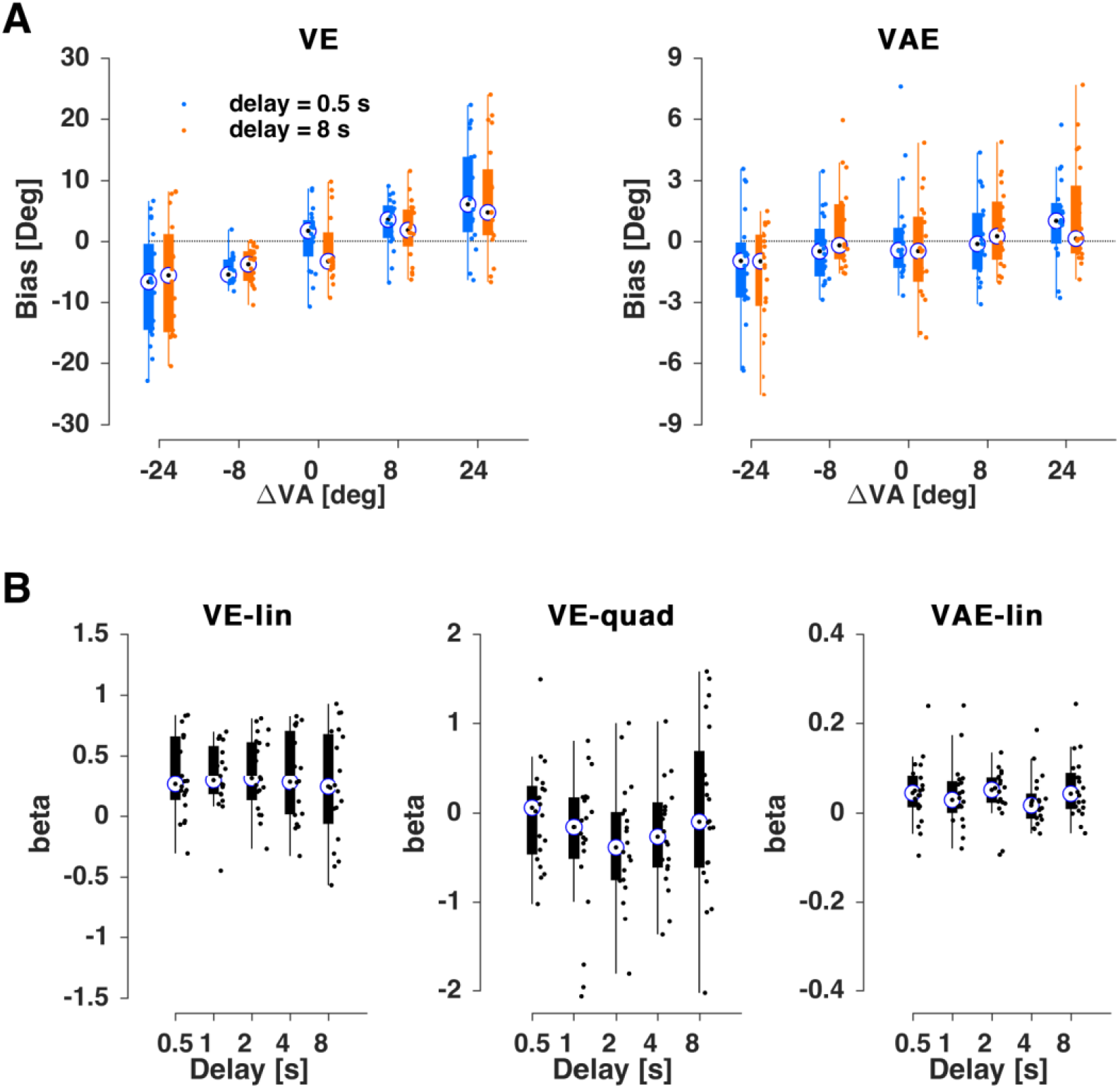
Results for experiment 1. **(A)** Ventriloquism bias (ve, left) and aftereffect bias (vae, right) for the shortest (0.5 s, blue) and longest (8 s, orange) delays. **(B)** Regression coefficients for participant-wise fits of the trial-averaged bias against the multisensory discrepancy (ΔVA). For the ve both linear (VE-lin) and non-linear (VE-quad) slopes are shown, for the aftereffect just the linear (VAE-lin). Boxplots indicate medians (bullet), quartiles, and individual data (dots).

We implemented two separate analyses to probe whether the biases differed as a function of delay. In a first approach, we fit a GLMM across all single trial biases, conditions and participants. Extending model 3 by the delay as an additional factor provided “very strong” evidence in favor of no effect of delay (BIC 55940 without and 55963 with including the delay; ΔBIC=23). The model parameters for the full model including the delay and its interactions revealed no significant effect for delay (**Table 1**).

**Table 1.** Generalized linear mixed-effects models for the ventriloquism effect (ve) and aftereffect (vae) in Experiment 1. **(A)** Model predicting the ve based on a linear and non-linear dependency on the multisensory discrepancy (ΔVA), including delay as factor (D). **(B)** Model predicting the vae based on a linear dependency on the multisensory discrepancy (ΔVA), including delay as factor (D). CI: 95% confidence interval (parametric).

| **(A) $ve \sim \beta_0 + \beta_1 \cdot \Delta VA + \beta_2 \cdot D + \beta_3 \cdot (\Delta VA)^{0.5} + \beta_4 \cdot \Delta VA:D + \beta_5 \cdot (\Delta VA)^{0.5}:D + (1/\text{subj})$** |  |  |  |  |  |
| --- | --- | --- | --- | --- | --- |
| Name | Estimate ($\beta$) | t-statistics | p-value | CI (95%) | |
| Intercept | -0.3527 | -0.6631 | 0.5073 | -1.3955 | 0.6900 |
| $\Delta VA$ | 0.0486 | 0.8784 | 0.3798 | -0.0598 | 0.1570 |
| D | -0.0291 | -0.6765 | 0.4988 | -0.1134 | 0.0552 |
| $(\Delta VA)^{0.5}$ | 1.1657 | 4.7161 | **0.0000** | 0.6812 | 1.6503 |
| $\Delta VA:D$ | 0.0042 | 0.3132 | 0.7541 | -0.0221 | 0.0304 |
| $(\Delta VA)^{0.5}:D$ | -0.0386 | -0.6460 | 0.5183 | -0.1559 | 0.0786 |

| **(B) $vae \sim \beta_0 + \beta_1 \cdot \Delta VA + \beta_2 \cdot D + \beta_3 \cdot \Delta VA:D + (1/\text{subj})$** |  |  |  |  |  |
| --- | --- | --- | --- | --- | --- |
| Name | Estimate ($\beta$) | t-statistics | p-value | CI (95%) | |
| Intercept | -0.0165 | -0.1177 | 0.9063 | -0.2909 | 0.2580 |
| $\Delta VA$ | 0.0377 | 4.2980 | **0.0000** | 0.0205 | 0.0549 |
| D | 0.0053 | 0.1564 | 0.8757 | -0.0612 | 0.0718 |
| $\Delta VA:D$ | 0.0012 | 0.5556 | 0.5785 | -0.0030 | 0.0053 |

In a second approach, we fit the participant and condition-wise trial-averaged biases using individual regression models and investigated whether these two slopes (linear, nonlinear) differed as a function of delay using a non-parametric test (**Figure 2B**): neither slope revealed an effect of delay (Friedman’s nonparametric ANOVA, reporting FDR corrected p-values; linear term: chi(4,99)=4.5, p_fdr_ =0.95, quadratic term: chi(4,99)=3.1, p_fdr_ =1.2). Hence, our data offer no evidence that manipulating the delay between the AV stimulus and the associated response affects the strength of the ventriloquism bias.

The same manipulation also did not affect the ventriloquism aftereffect (**Figure 2A**). The addition of the delay in model 1 resulted in a reduced fit (BIC without 52322 and with delay 52339; ΔBIC=17 providing “very strong” evidence in favor of no effect) and in the combined model neither the effect of delay nor its interaction with ΔVA were significant (**Table 1**). The analysis of participant-and condition-wise biases led to the same conclusion (**Figure 1C**; linear term: chi(4,99)=6.8, p_fdr_ = 0.6; **Figure 2B**).

### Manipulating the delay within auditory trials

In a second experiment (n=21) we tested whether adding a similar temporal delay between the auditory stimulus and the response in the A trial would affect the two biases (**Figure 3A**). First, and as expected given that the manipulation was specific to the A trial, we found that the ventriloquism bias was not affected: adding the delay as factor did not improve model fit (BIC without 57840 and with delay as factor 57865; ΔBIC=25 providing “very strong” evidence in favor of no effect) and the factor delay and its interactions were not significant (**Table 2**). The condition-and participant-wise biases were also not significantly different between delays (linear term: chi(4,104)=1.4, p_fdr_ = 1.7, quadratic term: chi(4,104)=1.6, p_fdr_ =1.7; **Figure 3B**).

**Table 2.**
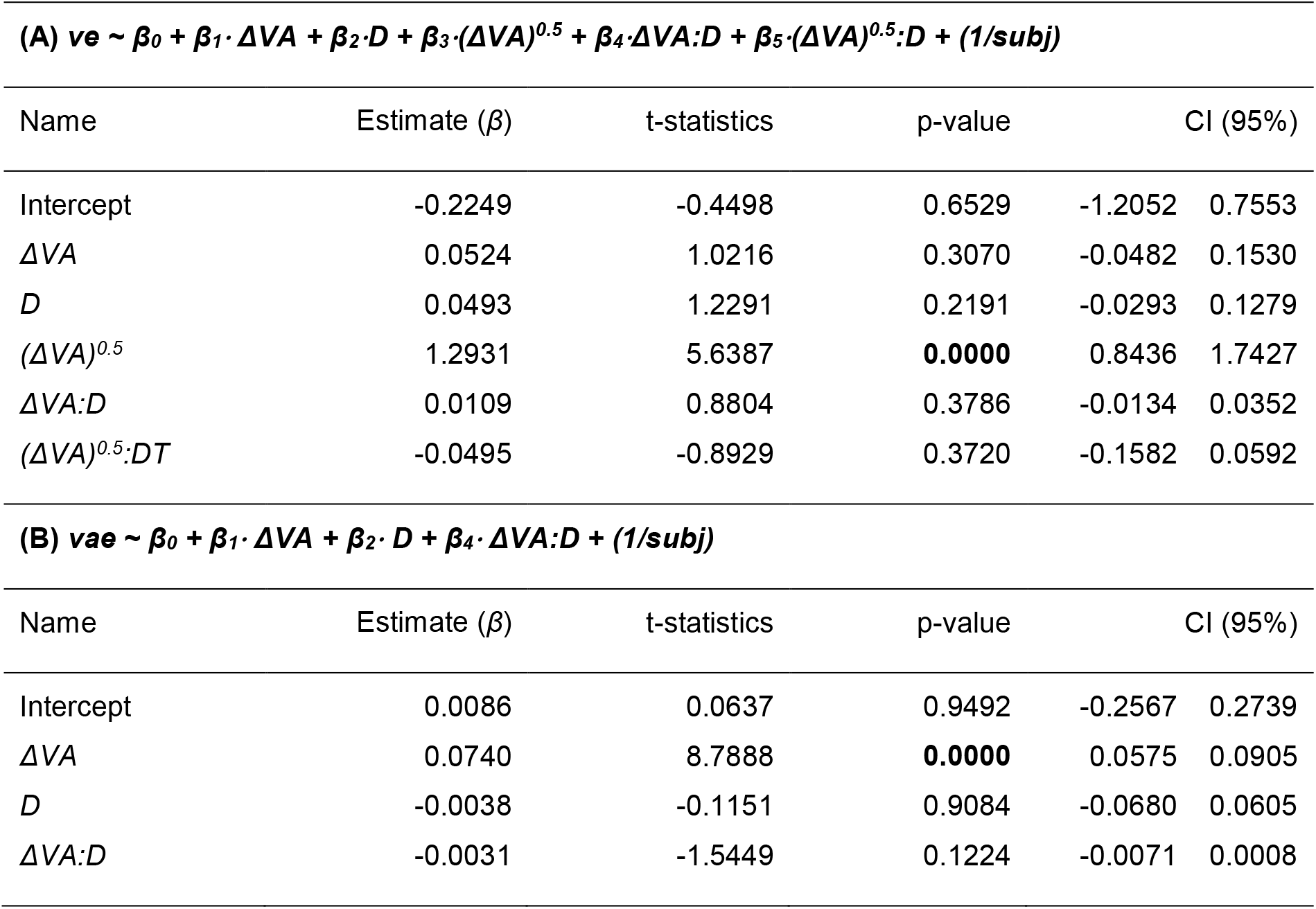
Generalized linear mixed-effects models for the ventriloquism effect (ve) and aftereffect (vae) for Experiment 2. **(A)** Model predicting the ve based on a linear and non-linear dependency on the multisensory discrepancy (ΔVA), including delay as factor (D). **(B)** Model predicting the vae based on a linear dependency on the multisensory discrepancy (ΔVA), including delay as factor (D). CI: 95% confidence interval (parametric).

**Figure 3.**
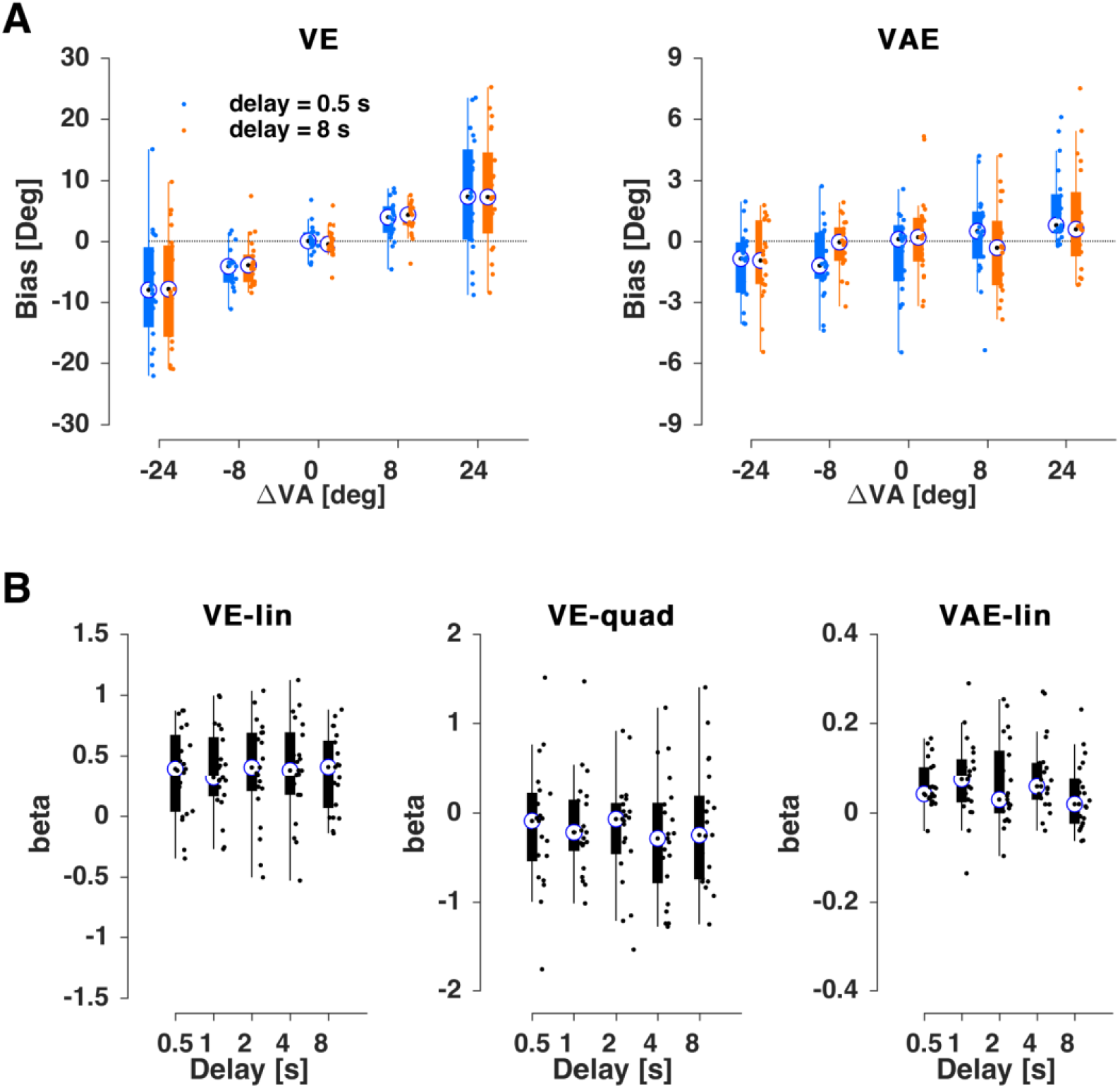
Results for Experiment 2. **(A)** Ventriloquism bias (ve, left) and aftereffect bias (vae, right) for the shortest (0.5 s, blue) and longest (8 s, orange) delays. **(B)** Regression coefficients for participant-wise fits of the trial-averaged bias against the multisensory discrepancy (ΔVA). For the ve both linear (VE-lin) and non-linear (VE-quad) slopes are shown, for the aftereffect just the linear (VAE-lin). Boxplots indicate medians (bullet), quartiles, and individual data (dots).

Interestingly, also the aftereffect did not change with the delay in this experiment: the addition of the delay did not improve the model fit (BIC 54617 vs. 54633; ΔBIC=15 providing “very strong” evidence in favor of no effect) and the interaction terms were not significant (**Table 2**). The same conclusion was supported by the participant-and condition-wise biases (linear term: chi(4,104)=8.1, p_fdr_ = 0.4; **Figure 3B**).

### Masking audio-visual trials

In a third experiment (n=22) we tested whether the ventriloquism aftereffect could be manipulated by adding a sensory-motor masker added in between the AV and A trials. The masker comprised a sensory component both in the visual (full-screen random dot pattern) and auditory modalities (a spatially diffuse sound) and required the participants to make a motor response to also mask potential memory traces of the preceding motor response in the AV trial. For comparison, participants performed blocks with the interleaved masking trial and without. We ensured that the overall temporal delay between the AV and A trials was comparable across these two conditions.

As in the two preceding experiments, we observed robust ventriloquism and aftereffect biases in the AV and A trials (**Figure 4A**). As expected given the experimental design, the ventriloquism effect did not differ significantly between conditions. The addition of the masking condition as factor did not improve model fit (BIC without 60978 and with masker 60998; ΔBIC=20 providing “very strong” evidence in favor of no effect) and the interaction terms were not significant (**Table 3**). The analysis of participant-wise biases confirmed this (linear slope: chi(1,43)=0.7, p_fdr_ = 0.8, quadratic slope: chi(1,43)=0.7, p_fdr_ =0.8; **Figure 4B**).

**Table 3.** Generalized linear mixed-effects models for the ventriloquism effect and aftereffect for Experiment 3 (masking effect in AV trial). **(A)** Model predicting the ve based on a linear and non-linear dependency on the multisensory discrepancy (ΔVA), including masking as factor (M). **(B)** Model predicting the vae based on a linear dependency on the multisensory discrepancy (ΔVA), including masking as factor (M). CI: 95% confidence interval (parametric).

| **(A) $ve \sim \beta_0 + \beta_1 \cdot \Delta VA + \beta_2 \cdot M + \beta_3 \cdot (\Delta VA)^{0.5} + \beta_4 \cdot \Delta VA:M + \beta_5 \cdot (\Delta VA)^{0.5}:M + (1/\text{subj})$** |  |  |  |  |  |
| --- | --- | --- | --- | --- | --- |
| Name | Estimate ($\beta$) | t-statistics | p-value | CI (95%) | |
| Intercept | 0.2768 | 0.3883 | 0.6978 | -1.1205 | 1.6741 |
| $\Delta VA$ | 0.2962 | 3.0450 | **0.0023** | 0.1055 | 0.4868 |
| $M$ | -0.1848 | -0.7841 | 0.4330 | -0.6469 | 0.2773 |
| $(\Delta VA)^{0.5}$ | 0.7435 | 1.5604 | 0.1187 | -0.1905 | 1.6775 |
| $\Delta VA:M$ | -0.0958 | -1.5484 | 0.1216 | -0.2171 | 0.0255 |
| $(\Delta VA)^{0.5}:M$ | 0.3473 | 1.1460 | 0.2518 | -0.2467 | 0.9413 |

| **(B) $vae \sim \beta_0 + \beta_1 \cdot \Delta VA + \beta_2 \cdot M + \beta_4 \cdot \Delta VA:M + (1/\text{subj})$** |  |  |  |  |  |
| --- | --- | --- | --- | --- | --- |
| Name | Estimate ($\beta$) | t-statistics | p-value | CI (95%) | |
| Intercept | 0.0013 | 0.0041 | 0.9967 | -0.6212 | 0.6239 |
| $\Delta VA$ | 0.0178 | 1.1577 | 0.2470 | -0.0123 | 0.0479 |
| $M$ | -0.0013 | -0.0065 | 0.9949 | -0.3972 | 0.3946 |
| $\Delta VA:M$ | 0.0159 | 1.6244 | 0.1043 | -0.0033 | 0.0351 |

**Figure 4.**
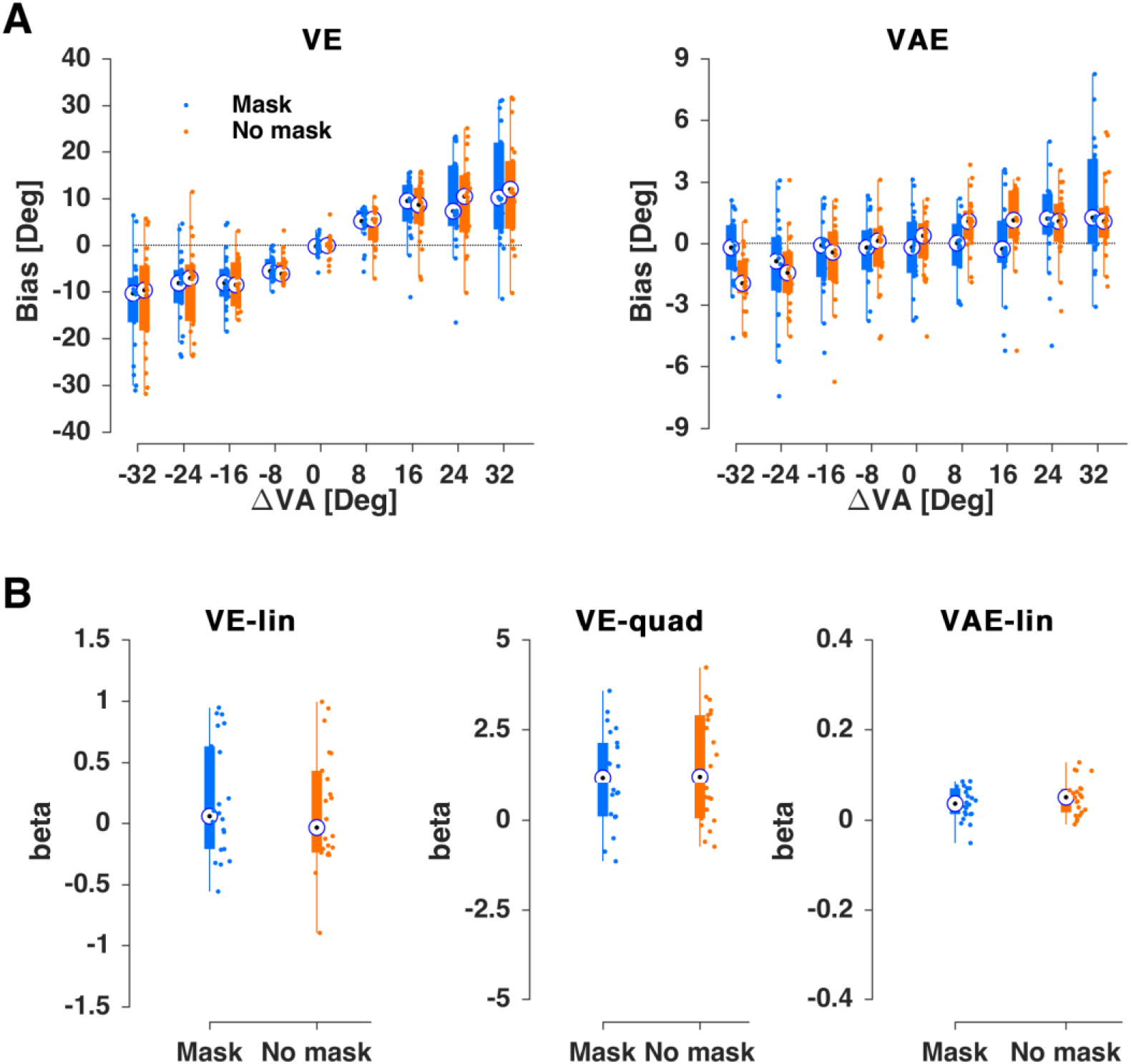
Results for Experiment 3. **(A)** Ventriloquism bias (ve, left) and aftereffect bias (vae, right) for the shortest (0.5 s, blue) and longest (8 s, orange) delays. **(B)** Regression coefficients for participant-wise fits of the trial-averaged bias against the multisensory discrepancy (ΔVA). For the ve both linear (VE-lin) and non-linear (VE-quad) slopes are shown, for the aftereffect just the linear (VAE-lin). Boxplots indicate medians (bullet), quartiles, and individual data (dots). Blue with masking trial; orange no masking trial.

Interestingly the masking manipulation did not affect the ventriloquism aftereffect. The addition of masking condition did not improve the model fit (BIC without 58439 and with delay 58455; ΔBIC=15 providing “very strong” evidence in favor of no effect) and the model parameters revealed no significant contribution of condition (**Table 3**). Finally, the analysis of individual participant data revealed no significant difference in slope (Fig. 4; chi(1,43)=1.6, p_fdr_ = 0.8; **Figure 4B**).

## DISSCUSSION

We tested the hypothesis that the trial-wise ventriloquism aftereffect is susceptible to manipulations known to interfere with working memory. Across three variations of an established ventriloquism paradigm we found no evidence for an interference of prolonged temporal delays (up to 8 seconds) or sensory-motor masking trials to reduce the strength of the ventriloquism aftereffect bias.

### The ventriloquism aftereffect and working memory

The motivation to probe the ventriloquism aftereffect against memory manipulations came from two observations. First, previous neuroimaging studies on the cerebral origin of the ventriloquism aftereffect have suggested a role of medial parietal regions such as the precuneus^7,15^. Studies on working memory or spatial navigation tasks have implied these regions in maintaining a persistent representation of multisensory spatial information^10,11,26,27^. We have previously shown that parietal representations of multisensory spatial information are maintained between trials in the ventriloquism paradigm, and are predictive of the aftereffect bias^7^. Hence, a role of short-term memory in the aftereffect is directly suggested by neuroimaging results.

Second, previous work on serial dependencies in unisensory perceptual tasks has suggested that these dependencies do not arise from sensory-level affects but rather reflect higher cognitive processes such as memory or the use of remembered information for subsequent decisions^16,18,22^. For example, a study on serial dependencies in visual judgements has used very similar mnemonic manipulations of temporal delays to show that the trial-wise biases are affected by the delay manipulation^22^. Hence, the observation that sensory and meta-cognitive variables carry over between trials even in simple laboratory paradigms also suggests a role of memory-related processes in the ventriloquism aftereffect.

While we did not find a dissipating effect of temporal delays or sensory maskers on the ventriloquism aftereffect, a previous study suggested that intervening audio-visual trials before the auditory trial lead to a reduction of the trial-wise aftereffect^4^. This suggests that multisensory information that bears the very same task-relevance can reduce the aftereffect, while a masking stimulus that comprises distinct audio-visual features and pertains to a different task does not, as seen in the present study. In addition, one study used repetitive AV trials to induce the ventriloquism aftereffect and found that this accumulates over repetitions but also dissipates over delays of 5 s and 20 s when no sensory interference is present^9^.

Hence, the combined evidence suggests that the trial-wise aftereffect and that induced by prolonged and repetitive exposure to a consistent audio-visual discrepancy differ in their sensitivity to memory interference. Still, future work is needed to directly test this hypothesis within the same participants and experimental design.

### Does the lack of evidence speak for the absence of an effect?

The absence of a significant result can naturally arise from a number of reasons. First, the sample size may have been too small. We based the sample size on general recommendations for behavioral tests^28^ and our previous studies^7,14^. Across several studies we found that a sample size of about 20 participants is sufficient to reliably detect both the ventriloquism effect and its aftereffect and the present data confirm this. Furthermore, the obtained effect sizes for the absence of an effect of delay or masking conditions (BIC differences) clearly speak against an effect rather than being inconclusive. Hence, the collective evidence obtained across the three experiments provides converging evidence that the ventriloquism aftereffect is robust against the tested manipulations.

It could also be that the tested memory manipulations did not interfere sufficiently with the relevant neural processes maintaining the sensory information. Longer delays may have a stronger influence^9^, but may came at the cost of overall reduced attention to the task, making it difficult to disentangle attention and memory effects. Here we restricted the maximal delay to 8 seconds to facilitate the collection of sufficiently many trials for all conditions within a single experimental session. Future studies could test alternative manipulations such as more extensive masking stimuli that provide a more comprehensive sensory-motor interference, or could consider the use of a dual task paradigm enhancing the simultaneous memory load.

### Implications for understanding the neural underpinnings of the ventriloquism aftereffect

Previous work has implied parietal regions and also early sensory regions in the ventriloquism aftereffect. For example, work on the long-term aftereffect suggested that the underlying processes involve the recalibration of early sensory representations, more so than relying only on high-level processes in fronto-parietal regions^15,29^. In contrast, in a recent study we found a primary role of parietal regions in mediating the trial-by-trial effect and contributing to long-term recalibration as well^7,14^. Combined with the behavioral results obtained here these neuroimaging studies suggest that the trial-wise aftereffect is not completely mediated by parietal regions involved in short-term memory, but rather originates from more distributed processes comprising regions that are insensitive to the present memory manipulations. One possibility for future work could be to directly quantify the maintenance of the audio-visual information received in the ventriloquist trial based on single-trial classification^14,14^ in order to probe the efficacy of the memory manipulations and to determine whether and where in the brain the trial-wise aftereffect is established despite memory interference.

## METHODS

We report data from three experiments, in which a sample of 20, 21 and 22 (not necessarily distinct) right-handed healthy young adults participated (age range: Exp 1: 19 – 30, mean±SD: 23.1±2.87; Exp 2: 18 – 30, mean±SD: 23.4±3.17 ; Exp 3: 20 - 30, mean±SD: 25.3 +-2.57). All had tested normal vision and reported normal hearing and no history of neurological or psychiatric disorders. Each participant provided written informed consent and was compensated monetarily. The study was approved by the local ethics committee of Bielefeld University.

### General experimental setup and task

The design of the experiments followed previous studies on the ventriloquism aftereffect^4,7^. Each of the three experiments was based on the same a single-trial localization task designed to probe both the audio-visual spatial ventriloquism effect and its aftereffect^4,7^. Participants sat 135 cm in front of an acoustically transparent screen (Screen International Modigliani, 2×1 m) with their head on a chin rest. Sounds were presented using a multi-channel soundcard (Creative Sound Blaster Z), amplified via an audio amplifier (t.amp E4-130, Thomann Germany) and played from one of five speakers (Monacor MKS-26/SW, MONACOR International GmbH, Germany) located behind the screen. The acoustic stimulus was a 1300 Hz sine wave tone (50 ms duration) sampled at 48 kHz and presented at 64 dB r.m.s. Visual stimuli were projected (Acer Predator Z650, Acer Inc., Taiwan) onto the screen. The visual stimulus was a cloud of white dots dispersed following a two dimensional Gaussian distribution (N = 200 dots, SD of vertical and horizontal spread 1.6°, width of a single dot = 0.12°, duration = 50 ms). Stimulus presentation was controlled using the Psychophysics toolbox (Brainard, 1997) for MATLAB (The MathWorks Inc., Natick, MA) with ensured temporal synchronization of auditory and visual stimuli.

Participants’ task was to localize a sound during either Audio-Visual (AV: sound and visual stimulus presented simultaneously) or Auditory (A: only sound) trials, or to localize a visual stimulus during Visual trials (V: only visual stimulus). Participants responded with a mouse cursor. Each trial started with a fixation period (Exp1,2: uniform 1100 ms – 1500 ms; Exp3: 1000 ms – 1200 ms) followed by the stimulus (50 ms). After a random post-stimulus period (see below) the response cue emerged, which was a horizontal bar along which participants could move a cursor. A letter ‘T’ was displayed on the cursor for ‘tone’ in the AV or A trials, and ‘V’ for the V trials to indicate which stimulus participants had to localize. There were no constraints on response times, however the participants were instructed to respond intuitively, and to not dwell too much on their response. Inter-trial intervals varied randomly (see below). A typical sequence of trials is depicted in **Figure 1**.

### Specific experimental designs

Experiments 1 and 2 manipulated the delay between the sensory stimulus and its respective response by inducing a variable delay (5 levels with mean delays of 0.5s, 1s, 2s, 4s, and 8s) between stimulus and the response cue. The precise delays were randomly jittered (uniform ± 200 ms) around these mean values to avoid participants forming specific expectations. Experiment 1 manipulated this delay for the AV trial, experiment 2 for the A trial. Each experiment consisted of 5 blocks, with each block comprising a sequence of 75 AV-A trials, and 15 interleaved V trials. For the AV trials, the locations of auditory and visual stimuli were drawn semi-independently from the 5 locations to yield a range of different audio-visual discrepancies (abbreviated ΔVA in the following; see below). For the A or V trials, stimulus locations were drawn from the 5 locations randomly. The audio-visual discrepancies in the AV trials took one of the following 5 values (ΔVA: -24°: -8°,0°,+8°,+24°), with the combinations of discrepancies and temporal delays changing pseudo-randomly across trials. Each combination of (ΔVA) and delay was repeated 15 times. Experiment 3 separated the AV and A trials by a sensory-motor masking trial. The masking trial consisted of an audio-visual display (duration for both audio and visual stimulus: 100 ms) and required participants to make a motor response. The visual mask was a more dispersed version of the standard visual stimulus with a SD of 5° (instead of 1.6°), centered at a location sample randomly from [-10°, 10°]. The auditory mask was the same standard sound stimulus but played from all 5 speakers and hence devoid of spatial information (**Figure 1B**, bottom). The motor masking task was to bring the cursor appearing randomly along the horizontal line to the middle red target box (**Figure 1B**, bottom). Experiment 3 comprised equal numbers of masking trials and no-masking (control) trials per level of ΔVA. Given that the stimulus and response in the masking trials required additional time, we extended the inter-trial interval between AV and A trials in the no-masking condition so that the average duration between the response in the preceding AV trial and the subsequent stimulus of the A trial was comparable between AV-A sequences with and without the masking trial (**Figure 1B**, bottom).

### Data analysis

The behavioral responses obtained in each trial were converted into response biases following previous studies^4^. The single-trial ventriloquism effect (ve) in the AV trials was defined as the difference between the actual sound location (A_AV_) and the reported sound location (R_AV_): ve = R_AV_ - A_AV_. The single-trial ventriloquism after-effect (vae) in the A trials was defined as the difference between the reported sound location (R_A_) and the mean reported location for all A trials of the same stimulus position (μR_A_), i.e., (vae = R_A_ - μR_A_), following previous work^4,7^.

Both biases are systematically related to the audio-visual discrepancy (ΔVA) in a linear, but possibly also in a nonlinear, manner^14,24,25^. For the ventriloquism bias the linear dependency describes the fusion of both stimuli for the response^24,30^, while the non-linear dependency describes the reduced tendency to bind multisensory stimuli when these are seemingly discrepant and not judged as arising from a common cause^24,30^. This nonlinear dependency on the ventriloquism bias on ΔVA follows Bayesian models of sensory causal inference and has been shown to better capture the behavioral bias in many ventriloquism-like paradigms compared to a pure linear model^24,31–33^. Given that the ventriloquism aftereffect is directly related to the sensory information received during the AV trial^6,8,34,35^, a similar linear and possibly nonlinear dependency is expected between the aftereffect bias and ΔVA. To determine the best dependency of each bias on ΔVA for the present dataset, we first compared three candidate models describing each bias. The respective GLMMs were fit across all single-trial biases (ve or vae) from all participants across all three experiments, regardless of the specific memory or masking manipulation:

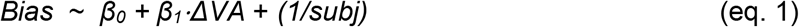

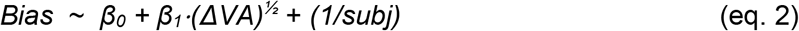

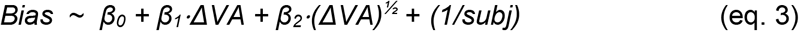

Here, and in the following, (ΔVA)^½^ stands for the signed square-root of the magnitude of ΔVA (i.e. sign(ΔVA) * sqrt(abs(ΔVA)), bias stands for the single-trial bias (ve, or vae), and subj stands for the participant id. The specific form of nonlinear dependency was chosen based on previous work^14,24,25^. Models were fit using a maximum likelihood procedure using the Laplace method in Matlab R2017a (fitglme.m). These models were compared based on their respective BIC. Interpretations of differences in BIC’s were based on established criteria, with values larger than 6 corresponding to “strong” and those larger than 10 to “very strong” evidence^36^. This revealed (see Results) that model 3 provided the best fit for the ventriloquism bias and model 1 for the aftereffect.

We then used two approaches to probe whether the ventriloquism bias or the aftereffect are significantly affected by the manipulations of the delays or the masking condition. Each experiment was analyzed separately in the following two ways. In a parametric approach, we extended the above models (model 1 for the vae; model 3 for the ve) by the trial-specific delay (in milliseconds) or the masking condition as additional (continuous or categorical) factors, including their interaction with the linear (and possibly also nonlinear) ΔVA-dependencies. Again we compared BIC values between the respective model without delay (masking condition) and the model including these. In addition, we investigated the respective model parameters and their confidence intervals (**Tables 1-3**).

In the second approach we asked whether the distribution of the condition-wise and participant-wise (trial-averaged) biases show a significant pattern indicative of an effect of delay (masking) manipulation. For this we modelled the trial-averaged participant-wise biases against a linear (vae) or combined linear and nonlinear (ve) dependency on ΔVA. We then used Friedmann’s non-parametric ANOVA to quantify whether the regression beta’s for the linear or nonlinear terms differed as a function of delay (masking condition). Here we corrected across multiple tests using the Benjamini & Yekutieli method for the false discovery rate (FDR)^37^.

## ACKNOWLEDGEMENTS

This work was supported by the European Research Council (to C.K. ERC-2014-CoG; grant No 646657). We would like to thank a number of undergraduate student and research assistants for their help during data collection.

## AUTHOR CONTRIBUTIONS

CK and HP Conceived study; collected and analyzed data; wrote the manuscript.

## ADDITIONAL INFORMATION

Authors declare no competing interests.

## REFERENCES

1. Chen, L. & Vroomen, J. Intersensory binding across space and time: A tutorial review. Atten Percept Psychophys 75, 790–811 (2013).

2. De Gelder, B. & Bertelson, P. Multisensory integration, perception and ecological validity. Trends in Cognitive Sciences 7, 460–467 (2003).

3. Alais, D. & Burr, D. The Ventriloquist Effect Results from Near-Optimal Bimodal Integration. Current Biology 14, 257–262 (2004).

4. Wozny, D. R. & Shams, L. Recalibration of Auditory Space following Milliseconds of Cross-Modal Discrepancy. Journal of Neuroscience 31, 4607–4612 (2011).

5. Recanzone, G. H. Rapidly induced auditory plasticity: The ventriloquism aftereffect. Proceedings of the National Academy of Sciences 95, 869–875 (1998).

6. Bruns, P. & Röder, B. Sensory recalibration integrates information from the immediate and the cumulative past. Sci Rep 5, 12739 (2015).

7. Park, H. & Kayser, C. Shared neural underpinnings of multisensory integration and trial-by-trial perceptual recalibration in humans. eLife 8, e47001 (2019).

8. Frissen, I., Vroomen, J. & de Gelder, B. The Aftereffects of Ventriloquism: The Time Course of the Visual Recalibration of Auditory Localization. Seeing and Perceiving 25, 1–14 (2012).

9. Bosen, A. K., Fleming, J. T., Allen, P. D., O’Neill, W. E. & Paige, G. D. Accumulation and decay of visual capture and the ventriloquism aftereffect caused by brief audio-visual disparities. Exp Brain Res 235, 585–595 (2017).

10. Martinkauppi, S. Working Memory of Auditory Localization. Cerebral Cortex 10, 889–898 (2000).

11. Curtis, C. E. Prefrontal and parietal contributions to spatial working memory. Neuroscience 139, 173–180 (2006).

12. Nyberg, L. & Eriksson, J. Working Memory: Maintenance, Updating, and the Realization of Intentions. Cold Spring Harb Perspect Biol 8, a021816 (2016).

13. Brodt, S. et al. Fast track to the neocortex: A memory engram in the posterior parietal cortex. Science 362, 1045–1048 (2018).

14. Park, H. & Kayser, C. The neurophysiological basis of short-and long-term ventriloquism aftereffects. http://biorxiv.org/lookup/doi/10.1101/2020.06.16.154161 (2020) doi: 10.1101/2020.06.16.154161.

15. Zierul, B., Röder, B., Tempelmann, C., Bruns, P. & Noesselt, T. The role of auditory cortex in the spatial ventriloquism aftereffect. NeuroImage 162, 257–268 (2017).

16. Talluri, B. C., Urai, A. E., Tsetsos, K., Usher, M. & Donner, T. H. Confirmation Bias through Selective Overweighting of Choice-Consistent Evidence. Current Biology 28, 3128-3135.e8 (2018).

17. Fritsche, M., Mostert, P. & de Lange, F. P. Opposite Effects of Recent History on Perception and Decision. Current Biology 27, 590–595 (2017).

18. Benwell, C. S. Y., Beyer, R., Wallington, F. & Ince, R. A. A. History biases reveal novel dissociations between perceptual and metacognitive decision-making. http://biorxiv.org/lookup/doi/10.1101/737999 (2019) doi: 10.1101/737999.

19. Kiyonaga, A., Scimeca, J. M., Bliss, D. P. & Whitney, D. Serial Dependence across Perception, Attention, and Memory. Trends in Cognitive Sciences 21, 493–497 (2017).

20. Farrell, S. et al. A test of interference versus decay in working memory: Varying distraction within lists in a complex span task. Journal of Memory and Language 90, 66–87 (2016).

21. Macoveanu, J., Klingberg, T. & Tegnér, J. Neuronal firing rates account for distractor effects on mnemonic accuracy in a visuo-spatial working memory task. Biol Cybern 96, 407–419 (2007).

22. Bliss, D. P., Sun, J. J. & D’Esposito, M. Serial dependence is absent at the time of perception but increases in visual working memory. Sci Rep 7, 14739 (2017).

23. Dekel, R. & Sagi, D. Perceptual bias is reduced with longer reaction times during visual discrimination. Commun Biol 3, 59 (2020).

24. Cao, Y., Summerfield, C., Park, H., Giordano, B. L. & Kayser, C. Causal Inference in the Multisensory Brain. Neuron 102, 1076-1087.e8 (2019).

25. Park, H., Nannt, J. & Kayser, C. Sensory-and memory-related drivers for altered ventriloquism effects and aftereffects in older adults. http://biorxiv.org/lookup/doi/10.1101/2020.02.12.945949 (2020) doi: 10.1101/2020.02.12.945949.

26. Crawford, L. E., Landy, D. & Salthouse, T. A. Spatial working memory capacity predicts bias in estimates of location. Journal of Experimental Psychology: Learning, Memory, and Cognition 42, 1434–1447 (2016).

27. Ester, E. F., Sprague, T. C. & Serences, J. T. Parietal and Frontal Cortex Encode Stimulus-Specific Mnemonic Representations during Visual Working Memory. Neuron 87, 893–905 (2015).

28. Simmons, J. P., Nelson, L. D. & Simonsohn, U. False-Positive Psychology: Undisclosed Flexibility in Data Collection and Analysis Allows Presenting Anything as Significant. Psychol Sci 22, 1359–1366 (2011).

29. Bruns, P., Liebnau, R. & Röder, B. Cross-Modal Training Induces Changes in Spatial Representations Early in the Auditory Processing Pathway. Psychol Sci 22, 1120–1126 (2011).

30. Rohe, T. & Noppeney, U. Reliability-Weighted Integration of Audiovisual Signals Can Be Modulated by Top-down Attention. eNeuro 5, ENEURO.0315-17.2018 (2018).

31. Rohe, T., Ehlis, A.-C. & Noppeney, U. The neural dynamics of hierarchical Bayesian causal inference in multisensory perception. Nat Commun 10, 1907 (2019).

32. Körding, K. P. et al. Causal Inference in Multisensory Perception. PLoS ONE 2, e943 (2007).

33. Wozny, D. R., Beierholm, U. R. & Shams, L. Probability Matching as a Computational Strategy Used in Perception. PLoS Comput Biol 6, e1000871 (2010).

34. Recanzone, G. H. Interactions of auditory and visual stimuli in space and time. Hearing Research 258, 89–99 (2009).

35. Radeau, M. & Bertelson, P. The After-Effects of Ventriloquism. Quarterly Journal of Experimental Psychology 26, 63–71 (1974).

36. Wagenmakers, E.-J. A practical solution to the pervasive problems of p values. Psychonomic Bulletin & Review 14, 779–804 (2007).

37. Benjamini, Y. & Yekutieli, D. The Control of the False Discovery Rate in Multiple Testing under Dependency. The Annals of Statistics 29, 1165–1188 (2001).

